# Cross-platform proteomics signatures of extreme old age

**DOI:** 10.1101/2024.04.10.588876

**Authors:** Eric R Reed, Kevin B Chandler, Prisma Lopez, Catherine E Costello, Stacy L Andersen, Thomas T Perls, Mengze Li, Harold Bae, Mette Soerensen, Stefano Monti, Paola Sebastiani

## Abstract

In previous work we used a Somalogic platform targeting approximately 5000 proteins to generate a serum protein signature of centenarians that we validated in independent studies that used the same technology. We set here to validate and possibly expand the results by profiling the serum proteome of a subset of individuals included in the original study using liquid chromatography tandem mass spectrometry (LC-MS/MS). Following pre-processing, the LC-MS/MS data provided quantification of 398 proteins, with only 266 proteins shared by both platforms. At 1% FDR statistical significance threshold, the analysis of LC-MS/MS data detected 44 proteins associated with extreme old age, including 23 of the original analysis. To identify proteins for which associations between expression and extreme-old age were conserved across platforms, we performed inter-study conservation testing of the 266 proteins quantified by both platforms using a method that accounts for the correlation between the results. From these tests, a total of 80 proteins reached 5% FDR statistical significance, and 26 of these proteins had concordant pattern of gene expression in whole blood. This signature of 80 proteins points to blood coagulation, IGF signaling, extracellular matrix (ECM) organization, and complement cascade as important pathways whose protein level changes provide evidence for age-related adjustments that distinguish centenarians from younger individuals.

## Introduction

Proteins in blood serum, cerebrospinal fluid, and urine have proven to be potent diagnostic and prognostic biomarkers of many diseases [1], in addition to their providing insights into the biological mechanisms underlying diseases. Progress in this area has relied on the increasingly sophisticated proteomics technology that has seen major advances in the past few years. In bottom-up analysis, the approach to liquid chromatography tandem mass spectrometry (LC-MS/MS) that has dominated the field for decades [2], proteins are first digested into peptides that are then separated by one or more chromatographic steps based on properties such as pI and hydrophobicity and analyzed via online mass spectrometry that produces both molecular weight (MS) and the MS/MS sequence information (MS2 and MS3) as the peptides elute. The large dynamic range of serum protein concentrations challenges LC-MS/MS-based serum proteomics workflows and therefore highly abundant proteins such as albumin are frequently depleted from samples prior to the analysis, increasing the complexity of sample preparation. Thus, this technology is still limited by the number of samples that can be analyzed simultaneously, the usual necessity for multistep sample preparation, the coverage that can be achieved, and the complexities of data processing [3].

LC-MS/MS has been the dominant technology for proteomics until the last decade that saw the emergence of high-throughput, reagent-based technologies from companies like Somalogic and Olink. The SomaScan technology developed by Somalogic [4] uses DNA-based aptamer reagents called somamers that bind to specific proteins in a sample, without the need for complex sample preparation and depletion of albumin in blood serum. The key innovation is to use the hybridization of somamers to the proteins present in the sample to convert the problem of measuring protein abundance into DNA sequencing of the reagents that can be done efficiently by using DNA arrays. The technology is high-throughput, and the latest platform includes reagents to detect more than 11,000 human proteins. The proximity-extension assay technology developed by Olink uses matched pairs antibodies labelled with oligonucleotides barcodes that bind to the proteins in a sample to measure protein abundance. The Somalogic technology has a much more comprehensive coverage than the proximity-extension assay technology developed by Olink that is limited by using antibodies, or mass spectrometry [5] based proteomics that is challenged by the wide dynamic range of protein abundances in the target. However, the specificity of many somamers is difficult to validate, and the concordance of proteomics results that use different approaches can be low [6, 7]. Unlike LC-MS/MS, the Somalogic and Olink techniques cannot provide the information necessary to identify novel proteins or to determine post-translational modifications. Thus, a combination of multiple proteomic technologies has advantages in terms of throughput, cost, and information content.

In the last few years, a variety of proteomics technologies have propelled the discovery of biomarkers of healthy aging and longevity [8–10], using LC-MS/MS [11] and Somalogic [8, 10, 12]. We used a Somalogic platform targeting approximately 5000 proteins to generate a serum protein signature of centenarians that we validated in independent studies that used the same technology [13]. We set out here to validate and possibly expand the results by profiling the serum proteome of a subset of individuals included in the original study using LC-MS/MS.

## Methods

### Samples

Proteomics profiling was performed on serum samples from blood obtained from 50 participants of the New England Centenarian Study (NECS) that included three age cohorts: centenarians, centenarians’ offspring, and subjects without familial longevity [14]. For mass spectrometry, the 50 samples were selected from the original pilot study of 224 subjects that had been previously profiled with SomaScan [15] to uniformly cover an age range 50 to 100 years (Table 1). The LC-MS/MS analyses used 10 tandem mass tags (TMT); samples were profiled in five pools of ten samples each (plus an 11^th^ channel containing the mixed 10 serum samples for normalization), and each pool was run in triplicate, resulting in a total of 150 sample profiles and 15 mixed sample profiles across 15 LC-MS/MS runs. Whole blood RNA sequencing transcriptomic and genotype data was obtained from the Long Life Family Study (LLFS), a family-based study of healthy aging and longevity [16]. These data include complete transcriptomic and genotypic profiles of 1,377 subjects. These subjects covered an age range of 24 to 107 years with a mean age of 69.1 years. Transcriptomic profiling was performed in 30 separate batches, with the number of subjects profiled per batch ranging from 23 to 82.

**Table 1:** Cohort subject counts in full dataset and 50-subject subset.

|  | Cohort |  |  |
| --- | --- | --- | --- |
|  | *Centenarian* | *Offspring* | *Control* |
| **Published Data** | 77 | 82 | 65 |
| **50 Sample Subset** | 9 | 17 | 24 |

### Mass Spectrometry Profiling

Full details of the sample preparation and the LC-MS/MS analyses are included in the supplemental material. Briefly, serum samples were subject to depletion of the top 12 most abundant serum proteins followed by trypsin/LysC digestion, and TMT labeling performed according to the manufacturer’s protocol. Peptide pools were analyzed on an Orbitrap Fusion Lumos Tribrid mass spectrometer (Thermo Scientific) interfaced to an M class nanoUPLC (Waters) via a TriVersa NanoMate nanoESI source (Advion). Peptide molecular weights were determined in the MS1 mode, and data dependent analyses were used to generate MS2 and MS3 spectra because acquisition of MS3 data minimizes interference from co-eluting components and thus increases the accuracy of quantification.

### Processing of Mass Spectrometry Data

Peptide quantification of raw mass spectrometry data was performed using MaxQuant 1.6.17 [17] using search parameters detailed in the supplemental material. A database consisting of reviewed protein sequences from the Uniprot *Homo sapiens* database, accession ID UP000005640, (downloaded Sept. 29, 2019), was used for all searches. Filtration criteria for protein matches included 1% false discovery rate, and ≥ 1 unique peptide resulting in a filtered set of 11,584 peptides across 1,473 proteins. Following peptide quantification, we removed 461 peptides associated with 12 depleted proteins with gene symbols: ALBU, APOA1, APOA2, CRP, A1AG1, A1AG2, A1AT, A2MG, FIB, HPT, IGH, TRFE. Next, we updated Uniprot IDs and mapped gene symbols using Uniprot’s “ID mapping” tool (https://www.uniprot.org/id-mapping) (performed on September 20, 2022), and removed a single peptide associated with Uniprot identifiers S4R460, which had been removed from the Uniprot database, resulting in 11,122 peptides assigned across 1,450 proteins.

We further filtered the peptides based on missingness, defined as having either a measured value of 0 or failure to be identified. First, for each peptide, we checked for association between missingness and age cohort using logistic regression adjusting for year-of-collection and gender that could suggest an informative missing data mechanism. To account for biological and technical variability, we used generalized estimating equations (GEE) using the geepack (v1.3.4) R package and a Bonferroni corrected p-value less than 0.05 for statistical significance. This analysis did not identify any associations between missingness and age cohort.

Next, we removed peptides with a high missingness rate, based on any of the following criteria:

- Missingness in at least 20% of profiles, i.e., 30 out of 150
- Missingness in at least 20% of batches, i.e. 3 out of 15
- Missingness in at least 20% of pools, i.e. 1 out of 5.

Of the 11,122 assigned peptides, 7,726 were removed based on high missingness criteria, resulting in 2,653 peptides across 398 proteins for subsequent analyses.

We obtained aggregated measurements of protein expression by summing measured values of peptides annotated to the same protein. Prior to the aggregation, missing peptide values were imputed by drawing from a uniform distribution with a range of 0 to the minimum peptide measurement of each batch. Each profile was then normalized by dividing their expression profiles by their respective 10% trimmed mean, followed by a log2-transformation. Finally, the normalized profiles were batch corrected to reduce the impact of technical variability using ComBat (v3.42.0) [18].

### Analysis of Mass Spectrometry Data

For mass spectrometry data, we evaluated the differences in the mean of the log2-protein expression between the three age cohorts –centenarians, centenarians’ offspring and subjects without familial longevity— using linear regression adjusting for year-of-collection and gender. We used GEE to account for within-sample variability of each triplicate, and we assessed the global differences between age cohorts using the log-likelihood ratio chi-square tests with 2 degrees of freedom. P-values were corrected for multiple hypothesis testing using the Benjamini-Hochberg False Discovery Rate (FDR) correction[19].

### Processing of SomaScan Data

The SomaScan data included in Sebastiani *et al.* (2021) [13] comprised 4,783 aptamers mapping to 4,116 proteins. We updated the aptamer protein annotations from SomaLogic version 3.0 to 4.1, removing 147 aptamers no longer included in more recent versions. Consistent with the mass spectrometry analysis, we updated Uniprot IDs and mapped gene symbols using Uniprot’s “ID mapping” tool (performed on September 20, 2022), further removing 233 aptamers mapping to mouse protein, Q99LC4, and updating an additional 36 aptamers. The filtered data comprised 4,403 aptamers in 3,887 proteins. Updated SomaScan aptamer assignments to Uniprot identifiers are given in Supplemental Table S1. A total of 266 proteins (353 aptamers) were shared across with the processed mass spectrometry data, 3,621 proteins (4,050 aptamers) were detected only in the SomaScan data, and 132 proteins were detected only in the mass spectrometry data.

### Analysis of SomaScan Data

We re-analyzed the subset of SomaScan data comprising the same 50 subjects profiled with mass spectrometry following the same procedure as the published SomaScan study [15]. Briefly, processed SomaScan measurements were first log2-transformed, and aptamer-specific outlier quantities were set to missing, based on values beyond three standard deviations of the 5% trimmed-mean. We next analyzed the differences in the mean of the log2-protein expression between centenarians, centenarians’ offspring and subjects without familial longevity using linear regression with the same model formulation and tested for global differences between age cohorts with ANOVA F-statistic testing on 2 and 45 degrees of freedom. Additionally, we re-performed multiple testing correction using the Benjamini-Hochberg False Discovery Rate (FDR) correction [19].

### Identification of conserved proteins associated with extreme old age

To identify proteins for which statistical associations between expression and age cohorts was conserved across the mass spectrometry and published SomaScan studies we applied the adjusted maximum p-value conserved (AdjMaxP) association method to pairs of differential results of proteins shared across the mass spectrometry-based and SomaScan platforms [20]. Briefly, this method aggregates the nominal p-values of statistical results for shared features (proteins in this case) across studies to a single statistical test based on the maximum p-value from each feature-level series of tests, while accounting for inter-study dependencies arising from shared samples across studies. In cases where multiple aptamers were annotated to the same protein, we allowed individual mass spectrometry proteins to be paired with each multiple SomaScan aptamers result, resulting in 353 shared feature pairs. Through the evaluation of the inter-study dependence, the mass spectrometry and published SomaScan of these shared features constituted 1.77 effective studies rather than 2.00 studies if the data were independent. For multiple hypothesis correction, we considered a total of 4,535 features, including the 132 proteins that were only presented in the mass spectrometry data, 4,050 aptamers that were annotated to proteins available only in the SomaScan data, and the 353 shared feature pairs. The nominal p-values from the respective study of these 4,535 features were included in FDR correction of the 353 AdjMaxP p-values.

We further evaluated their conservation across platforms based on the consistency of directions of differences between centenarians and either offspring or controls, with fully conserved proteins demonstrating consistency of both comparisons between centenarians and control and centenarians and offspring, and partially conserved proteins demonstrating consistency for only one of the two comparisons. For partial conservation, the direction of effect was assigned based on the conserved comparison.

### RNA sequencing profiling

Total RNA was extracted from PAXgene tubes using the PAXgene blood miRNA kit (Qiagen Inc.) on the QIAcube (Qiagen Inc.). RNA concentration and integrity were assessed using the Agilent 4200 Tapestation. At the McDonnell Genome Institute (MGI) at Washington University, analysis against the sequencing library was performed with 500 ng to 1 μg of total RNA. Ribosomal RNA was blocked using FastSelect reagents (Qiagen Inc.) during cDNA synthesis. RNA was fragmented in reverse transcriptase buffer with FastSelect reagent and heating to 94 degrees for 5 minutes, 75 degrees for 2 minutes, 70 degrees for 2 minutes, 65 degrees for 2 minutes, 60 degrees for 2 minutes, 55 degrees for 2 minutes, 37 degrees for 5 minutes, 25 degrees for 5 minutes. mRNA was reverse transcribed to yield cDNA using SuperScript III RT enzyme (Life Technologies, per manufacturer’s instructions) and random hexamers. A second strand reaction was performed to yield ds-cDNA. cDNA was blunt ended, had an A base added to the 3’ ends, and then had Illumina sequ’ncing adapters ligated to the ends. Ligated fragments were then amplified for 15 cycles using primers incorporating unique dual index tags. Fragments were sequenced on an Illumina NovaSeq-6000 using paired end reads extending 150 bases. Basecalls and demultiplexing were performed with Illumina’s bcl2fastq software and a custom Python demultiplexing program with a maximum of one mismatch in the indexing read. After sequencing, reads were aligned to the human genome sequence GRCh38 with GENCODE annotations by using STAR [21].

### Processing of transcriptomic data

The LLFS transcriptomic profiling data included 1,377 individuals aged between 24 and 107 years and 60,649 transcripts. We removed low quality samples, based on intergenic reads percentage > 8% and possible samples swap based on gender mismatch, resulting in the removal of 29 profiles. Raw read counts were then normalized using DESeq2 [22], followed by log2-transformation. Finally, we removed lowly expressed transcripts with at least 10 counts per million in fewer than 3% of samples. The final filtered data set comprised 1,348 subjects and 11,173 genes.

### Analysis of transcriptomic data

We examined the effect of age at blood draw on each transcript levels by using a linear mixed-effect model, in which the transcript data was the dependent variable, age was the main predictor, and additional covariates included gender, education level, enrollment site, sequencing batch, percentage of intergenic reads, and the first four genome-wide principal components calculated from genetic data to adjust for genetic ancestry. To account for relatedness, the model included a random intercept with variance covariance matrix proportional to the genetic relation matrix. Genome-wide principal components and the genetic relation matrix were estimated from whole genome sequence data using the R/3.6.0 packages PC-Relate and PC-Air following the method by Conomos et al [23], using the GENESIS R package (v2.6.0) [24].Full details of the genetic data and the modeling approach are reported by Gurinovich et al [25]. Modelling was performed on 1,346 individuals with complete transcriptome, genotype, and covariate data. P-values were corrected for multiple hypothesis testing using the Benjamini-Hochberg (FDR) correction [19].

### Functional analysis of protein signatures

We performed functional analysis of protein signatures with hypergeometric test-based enrichment analysis of functionally annotated protein sets, as well as annotation of protein-protein interactions. Enrichment analysis was performed using the hypeR R package (v1.10.0) [26] using as background the total number of proteins across the mass spectrometry and SomaScan data, i.e. 4,019 proteins. Signatures were tested for over-representation of protein sets from Gene Ontology Molecular Function [27, 28] and Reactome [29] obtained from the mSigDB (v7.5.1)[30]. Prior to running hypeR, mSigDB gene sets were converted to Uniprot identifiers using Uniprot’s “ID mapping” tool (performed on October 15, 2022). Annotation of protein-protein interactions was performed by querying Uniprot identifiers with STRING database (v11.4) [31].

## Results

### Cross platform signatures of extreme old age-associated proteins

In our analyses, we sought to identify serum proteins that were associated with extreme old age in the two proteomics platforms: mass spectrometry and SomaScan array. Our data included serum protein profiles of 50 NECS participants that were measured by both platforms, and our analysis evaluated global differences in the mean expression between three age-related cohorts: centenarians, centenarians’ offspring (offspring) and subjects without familial longevity (controls), comprised of 9, 17, and 24 subjects (Table 1), respectively. The SomaScan data of 50 NECS participants was a subset of the 224 NECS participants profiles included in the proteomics signature of extreme old age previously reported [13]. Following pre-processing, the mass spectrometry data quantified 398 proteins, the SomaScan data included 3,887 proteins measured by 4,403 aptamers, with 266 proteins shared by both platforms linked to 353 SomaScan aptamers. The complete list of proteins, including Uniprot identifiers, SomaScan aptamer identifiers, protein symbols, and gene symbols are in Supplemental Table S2.

At 1% FDR statistical significance threshold, our analysis detected only 1 protein, IGFBP2, associated with extreme old age in the reduced SomaScan data set, and 44 proteins, including IGFBP2, in the mass spectrometry data set (Figure 1A, Table 2, Supplemental Table S2). In contrast, following update of aptamer-to-protein annotations, the analysis of the complete SomaScan study with 224 subjects discovered 1,229 proteins. Of the 266 proteins quantified by both platforms, 33 and 106 proteins were discovered as extreme old age markers in the mass spectrometry study and the published SomaScan study, respectively, and 23 proteins were discovered across both platforms, representing a generally high congruence as indicated by Fisher’s exact test, p-value = 0.0021 (Figure 1B, Table 2). Accordingly, of the 266 proteins quantified by both platforms, 10 and 83 were only discovered by mass spectrometry or the published SomaScan study, respectively.

**Table 2:**
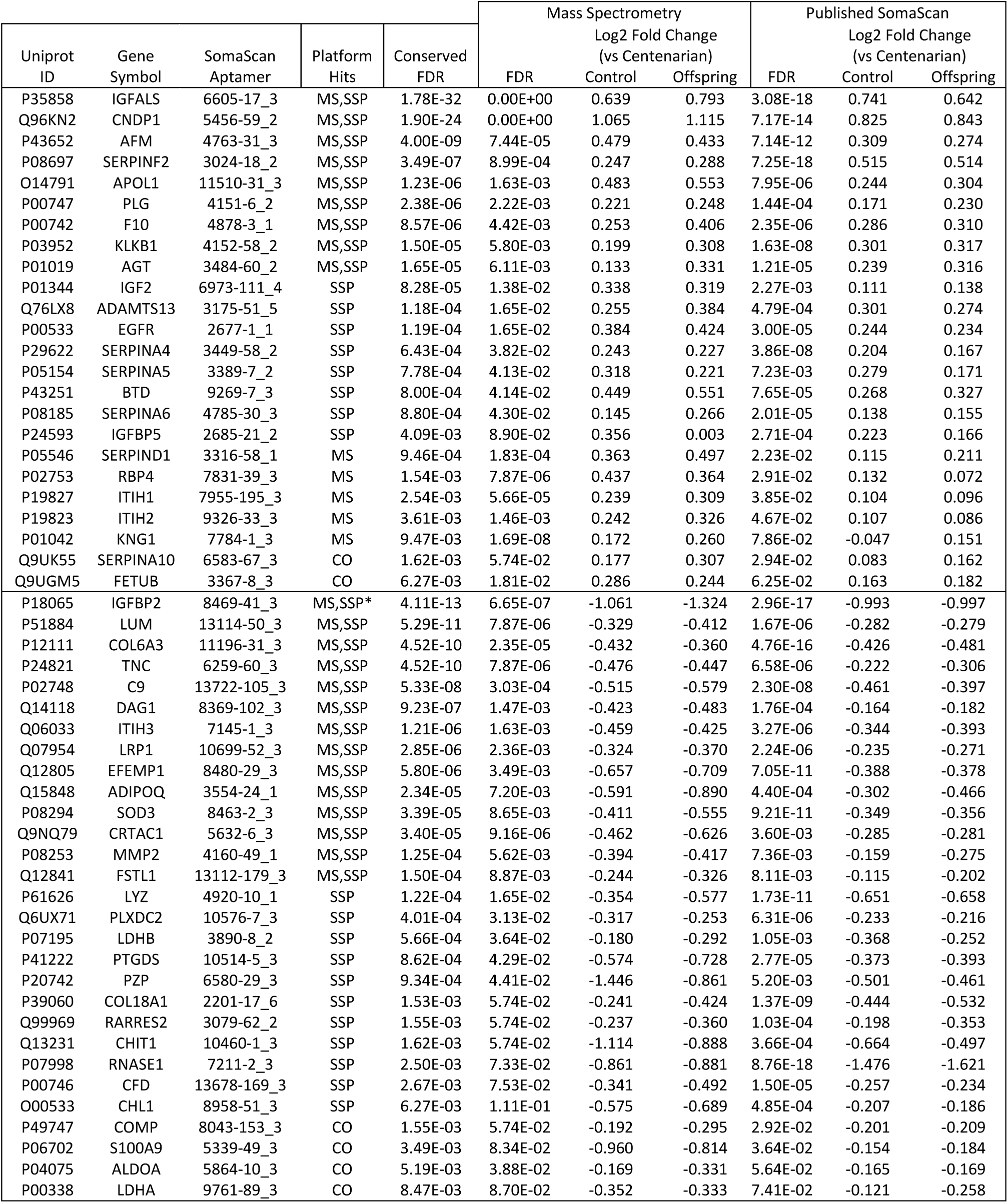
Inter-study validated protein signature (FDR < 0.01). The top half of the table shows proteins that are decreasing with older age, while the bottom half shows protein increasing at older age. Platform hits: SSP=only SomaScan; MS= only mass spectrometry; MS,SSP= both; CO=Conserved only * Also identified from the 50-sample subset of SomaScan data

**Figure 1:**
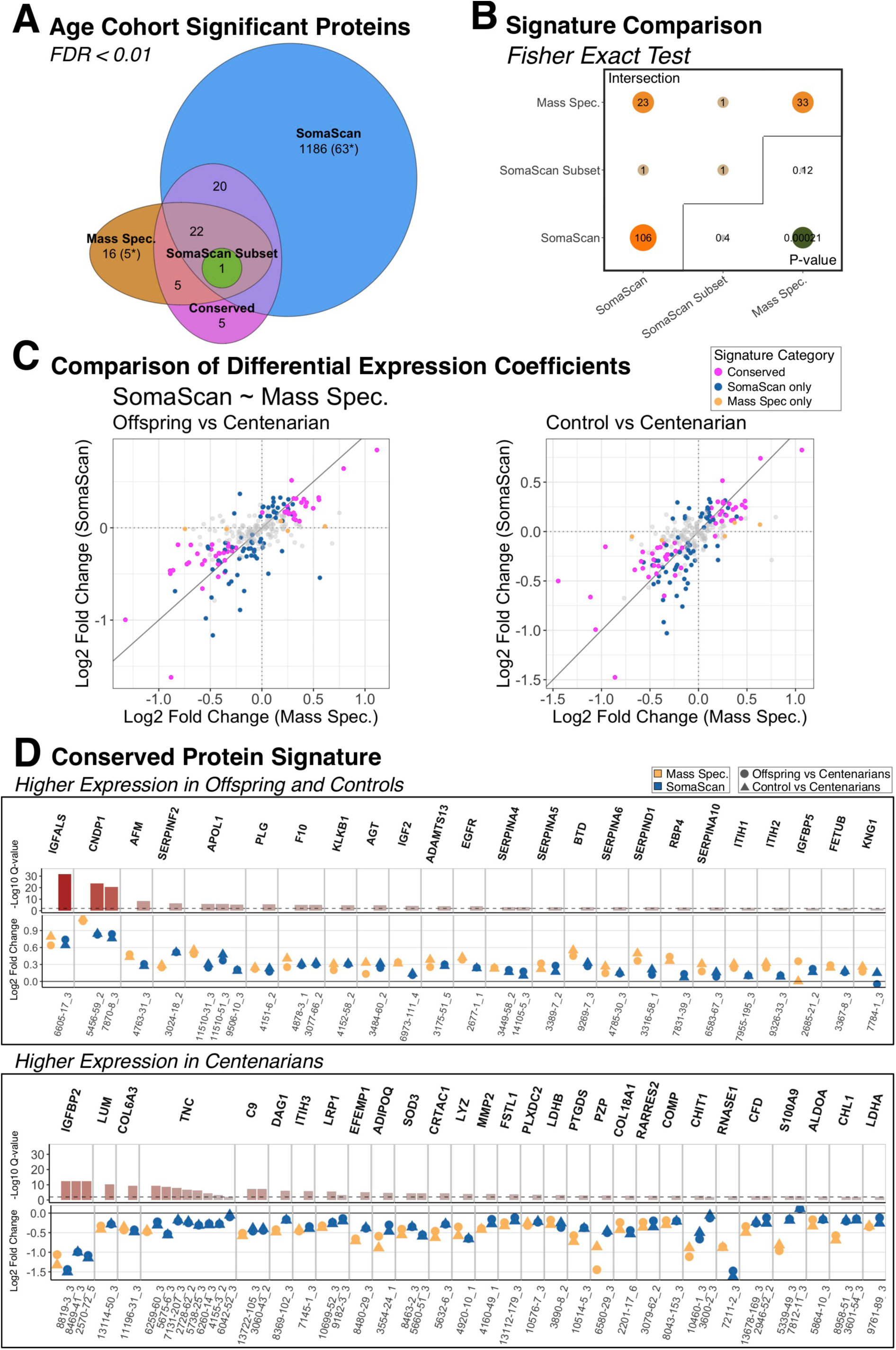
Inter-study protein signatures comparing centenarians to offspring and unrelated controls. A. Euler plot of protein signatures detected in the mass spectrometry data set with 50 subjects (Mass Spec), the SomaScan subsetwith the same 50 subjects (SomaScan Subset), and the SomaScan data with 224 participants (Somascan). Numbers in parentheses with a “*” denote the proteins that were included in both data sets but detected only in one analysis. B. Fisher’s Exact Test results of the intersection between mass spectrometry and SomaScan signatures. The diagonal represents the number of significant proteins from each analysis that are annotated by the SomaScan assay. C. Comparison of log2 fold changes protein expression differences of controls and offspring to centenarians from mass spectrometry and SomaScan studies across their 266 shared proteins. D. Log2 fold changes protein expression differences inter-study conserved proteins, including each mass spectrometry result and SomaScan aptamer. All protein signature results shown were identified using an FDR cutoff of 0.01. The full set of analysis results are reported in Supplementary Table S2.

To identify proteins for which associations between expression and age-cohorts were conserved across the mass spectrometry and the published SomaScan results, we performed inter-study conservation testing of the 266 proteins quantified by both platforms using the AdjMaxP method. From these tests, a total of 53 proteins reached 1% FDR statistical significance (Table 2, Supplemental Table S2). Of these 53 proteins, 52 demonstrated full cross-platform conservation based on consistent directions of differences between centenarians and either offspring or controls, and 1 protein, KNG1, demonstrated partial conservation due to consistently higher expression in offspring compared to centenarians, but we observed discrepancy in direction of difference between controls and centenarians (Figure 2C, Table 2). This set of 53 fully or partially conserved proteins included 25 proteins that had been previously identified by only one study at 1% FDR statistical significance, including 5 and 20 proteins that had been discovered by mass spectrometry or the published SomaScan study, respectively, as well as 5 proteins that were not previously identified by either study (Figure 1A, Table 2). Moreover, of these 53 proteins, 24 and 29 proteins were assigned as having higher expression in offspring/controls or centenarians, respectively (Figure 2D).

**Figure 2:**
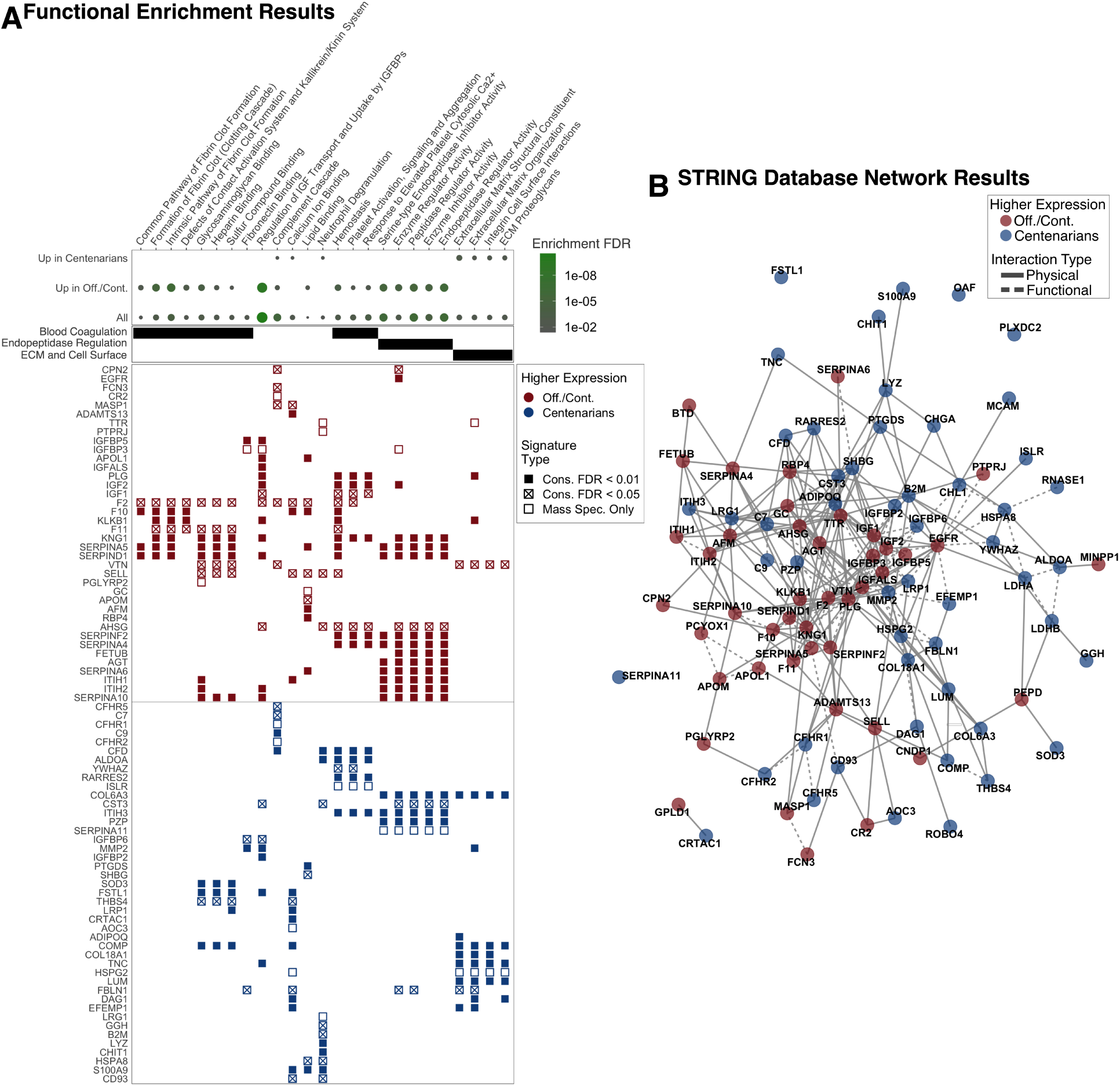
Protein interaction and functional analysis results. These results reflect analyses of 96 protein signature of proteins with conserved (FDR values < 0.05) and mass spectrometry-only discovered proteins (FDR < 0.01). A. Overrepresentation-based enrichment analysis comparing Gene Ontology Functional Terms and Reactome pathways to three protein lists: All 96 proteins (All), 44 proteins with higher expression in offspring and controls (Up in Off./Cont.), and 52 proteins with higher expression in centenarians (Up in Centenarians). Additional information for these results, including term source, p-values, set sizes, and set members are reported in Supplementary Table S3. B. STRING database annotated protein physical and functional interactions. The full list of interaction pairs is reported in Supplementary Table S4.

At 5% FDR threshold, an additional of 30 proteins reached statistical significance. However, three of these proteins, TGFBI, GAPDH, and DPEP2, did not have consistent patterns of associations across platforms (Supplemental Table S2). Of the remaining 27 proteins, 25 demonstrated full conservation, and two, F2 and CPN2, demonstrated partial conservation with consistent higher expression in either controls or offspring, respectively (Table 3). This additional set of 27 fully or partially conserved proteins included 18 proteins that had been previously identified by only one study at 1% FDR statistical significance, all of which had been discovered by the published SomaScan study, as well as 9 proteins that were not previously identified by either study (Table 3). Moreover, of these 27 proteins, 12 and 15 were assigned as having higher expression in offspring/controls or centenarians, respectively.

**Table 3:**
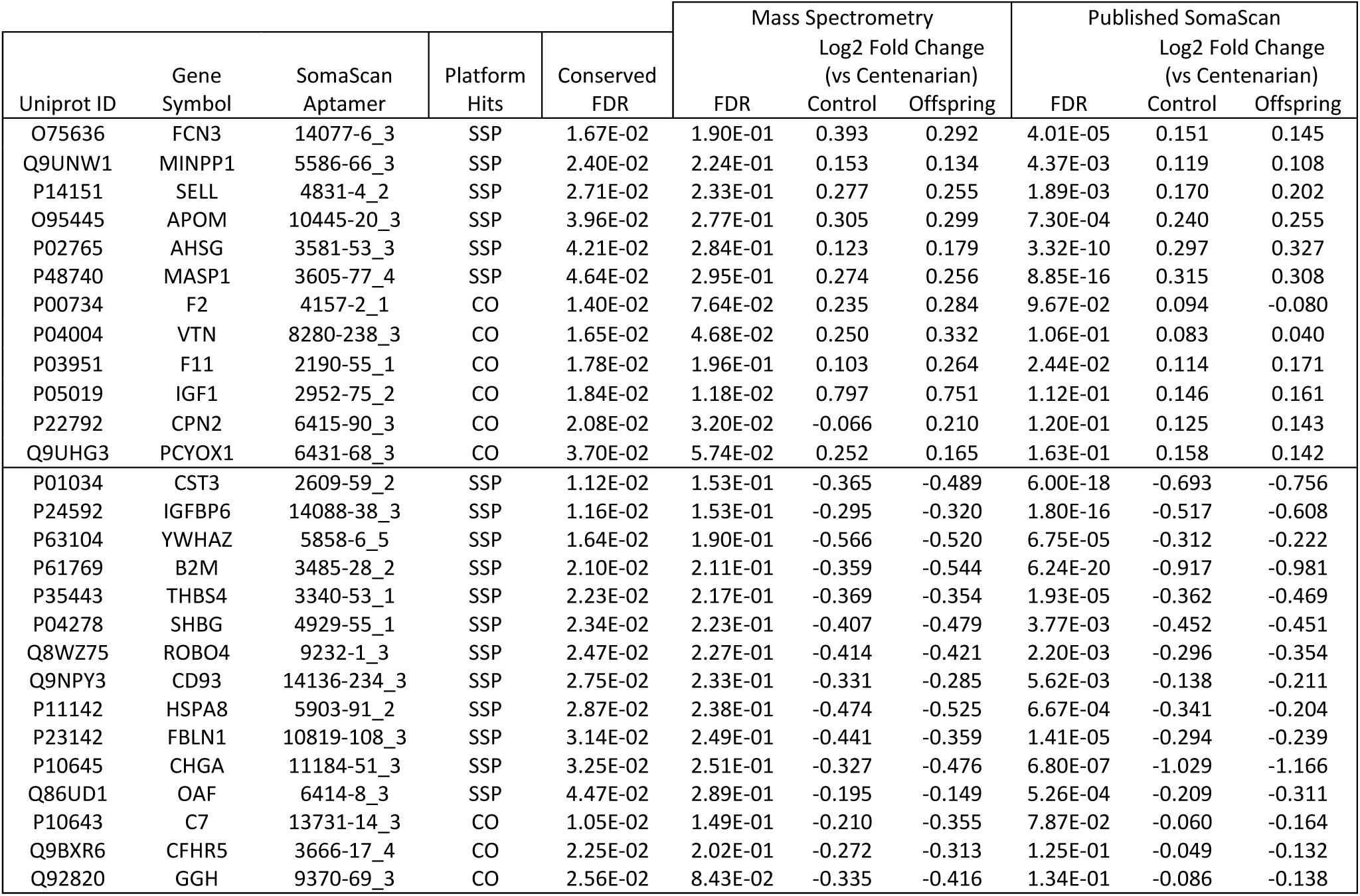
Additional inter-study validated protein signature (FDR < 0.05)

Finally, the mass spectrometry study identified 16 proteins at 1% FDR statistical significance threshold, 11 of which were either not measured by SomaScan and 5 proteins that were measured by SomaScan but were not identified by inter-study conservation tests. These 5 proteins that were not measured by SomaScan included IGFBP3, PGLYRP, PTPRJ, CFHR1, and SERPINA11 (Figure 1A, Table 4). Of these 16 proteins, 8 and 8 were assigned as having higher expression in offspring/controls or centenarians, respectively. Given that these 16 proteins were not identified in the published SomaScan study [13], either through lack of aptamer targets or failing to reach statistical significance, we included them in downstream analyses.

**Table 4:** Mass spectrometry only protein signature.

| Uniprot ID | Gene Symbol | SomaScan Aptamer | Platform Hits | Conserved FDR | Mass Spectrometry |  |  | Published SomaScan |  |  |
| --- | --- | --- | --- | --- | --- | --- | --- | --- | --- | --- |
|  |  |  |  |  | FDR | Log2 Fold Change (vs Centenarian) |  | FDR | Log2 Fold Change (vs Centenarian) |  |
|  |  |  |  |  |  | Control | Offspring |  | Control | Offspring |
| P17936 | IGFBP3 | 2571-12_3 | MS | 2.46E-01 | 8.59E-07 | 0.634 | 0.612 | 4.49E-01 | 0.070 | 0.016 |
| Q96PD5 | PGLYRP2 | 5601-2_3 | MS | 8.64E-02 | 1.98E-03 | 0.373 | 0.184 | 2.59E-01 | 0.090 | 0.072 |
| Q12913 | PTPRJ | 8250-2_3 | MS | 2.70E-01 | 6.90E-03 | 0.271 | 0.251 | 4.74E-01 | -0.048 | -0.033 |
| P80108 | GPLD1 |  | MS |  | 6.19E-13 | 0.662 | 0.624 |  |  |  |
| P02766 | TTR |  | MS |  | 1.40E-05 | 0.751 | 0.685 |  |  |  |
| P20023 | CR2 |  | MS |  | 2.22E-03 | 0.640 | 0.397 |  |  |  |
| P02774 | GC |  | MS |  | 2.46E-03 | 0.144 | 0.244 |  |  |  |
| P12955 | PEPD |  | MS |  | 9.92E-03 | 0.241 | 0.314 |  |  |  |
| Q03591 | CFHR1 | 5982-50_3 | MS | 4.24E-01 | 3.16E-04 | -0.694 | -0.757 | 6.11E-01 | -0.051 | -0.012 |
| Q86U17 | SERPINA11 | 9002-36_3 | MS | 7.26E-02 | 8.30E-03 | -0.363 | -0.333 | 2.35E-01 | -0.088 | -0.014 |
| P36980 | CFHR2 |  | MS |  | 6.49E-05 | -0.941 | -0.997 |  |  |  |
| O14498 | ISLR |  | MS |  | 9.37E-05 | -0.194 | -0.355 |  |  |  |
| P98160 | HSPG2 |  | MS |  | 8.56E-04 | -0.251 | -0.232 |  |  |  |
| P02750 | LRG1 |  | MS |  | 1.26E-03 | -0.568 | -0.554 |  |  |  |
| Q16853 | AOC3 |  | MS |  | 1.88E-03 | -0.285 | -0.462 |  |  |  |
| P43121 | MCAM |  | MS |  | 6.41E-03 | -0.346 | -0.225 |  |  |  |

### Functional analysis results

We performed functional annotation of 96 proteins, comprising the 80 proteins identified as associated with extreme old age by inter-study conservation analysis, and 16 additional proteins identified only by mass spectrometry using overrepresentation-based enrichment analysis. The compendium used for annotation including the Gene Ontology Molecular Function [27, 28] and the Reactome [29] knowledge bases (Figure 2A, Supplemental Table S3). Although most highly enriched categories included proteins assigned as having higher expression in either offspring/controls or centenarians, the enrichment of these categories was generally specific to either signature. The complement cascade was the only pathway that showed differential enrichment in both signatures and included six proteins with higher expression in offspring/controls, FCN3, MASP1, VTN, F2, CPN2, CR2, and six proteins with higher expression in centenarians, C9, CFD, CFHR1, C7, CFHR5, CFHR2.

Proteins with higher expression in offspring/controls were specifically enriched for functional categories related to insulin-like growth factor (IGF) signaling regulation, blood coagulation pathways, and endopeptidase regulation. Regulation of IGF transport and uptake by insulin-like growth factor binding proteins (IGFBPs) was the most significantly enriched category across all tests, FDR = 3.00E-11, and included 14 proteins assigned to this set, highlighted by the presence of IGFs, IGF1 and IGF2, IGFBs, IGFBP3 and IGFBP5, and IGFBP acid labile subunit (IGFALS). Moreover, two additional proteins, IGFBP2 and IGFBP6, demonstrated higher expression in centenarians. Notably, IGFALS and IGFBP2, were the most statistically significant proteins identified by cross-platform conservation analysis assigned to either cohort set, FDR = 1.78E-32 and 4.11E-13, respectively (Table S2, Figure 1C). Blood coagulation pathway results spanned eight functional categories, including four fibrin clot and contact activation system, and three platelet activity pathways. Finally, endopeptidase regulator activity related pathways spanned five functional categories, and included 12 proteins with higher expression in offspring/controls, highlighted by six serine-type endopeptidase inhibitor (SERPIN) proteins, SERPINF2, SERPINA4, SERPINA5, SERPINA6, SERPIND1, and SERPINA10.

Proteins with higher expression in centenarians were specifically enriched for categories related to extracellular matrix (ECM), cell surface, and neutrophil degranulation. Combined, this signature includes ten cell surface proteins, including CHL1, MCAM, PLXDC2, B2M, DAG1, LRP1, C7, C9, CD93, and ROBO4, and six extracellular matrix proteins, including COMP, TNC, LUM, HSPG2, COL6A3, COL18A1. Finally, the neutrophil degranulation category included 11 proteins: CHIT1, CST3, LRG1, GGH, LYZ, ALDOA, B2M, CD93, CFD, and HSPA8. (In some cases, the same protein appears in more than one of these categories.) Increased activity of genes involved in neutrophil degranulation is consistent with the shift from noncytotoxic to cytotoxic immune cells that we observed in in centenarians [32].

Given the extent to which these signature genes reside in functional categories, we sought to further explore their functional and physical interactions using the STRING database [31]. The database reported a total of 306 total interactions, and 78 physical interactions above at a confidence level above 0.4 (Figure 2B, Supplemental Table S4). To gain further insight into the STRING reported interactions, as well additional unreported interactions, we performed additional interrogation via Reactome [29] and literature review. This resulted in the characterization of 53 interactions, delineated into four categories, binding, activation, inhibition, and proteolysis (Figure 3, Supplemental Table S5).

**Figure 3:**
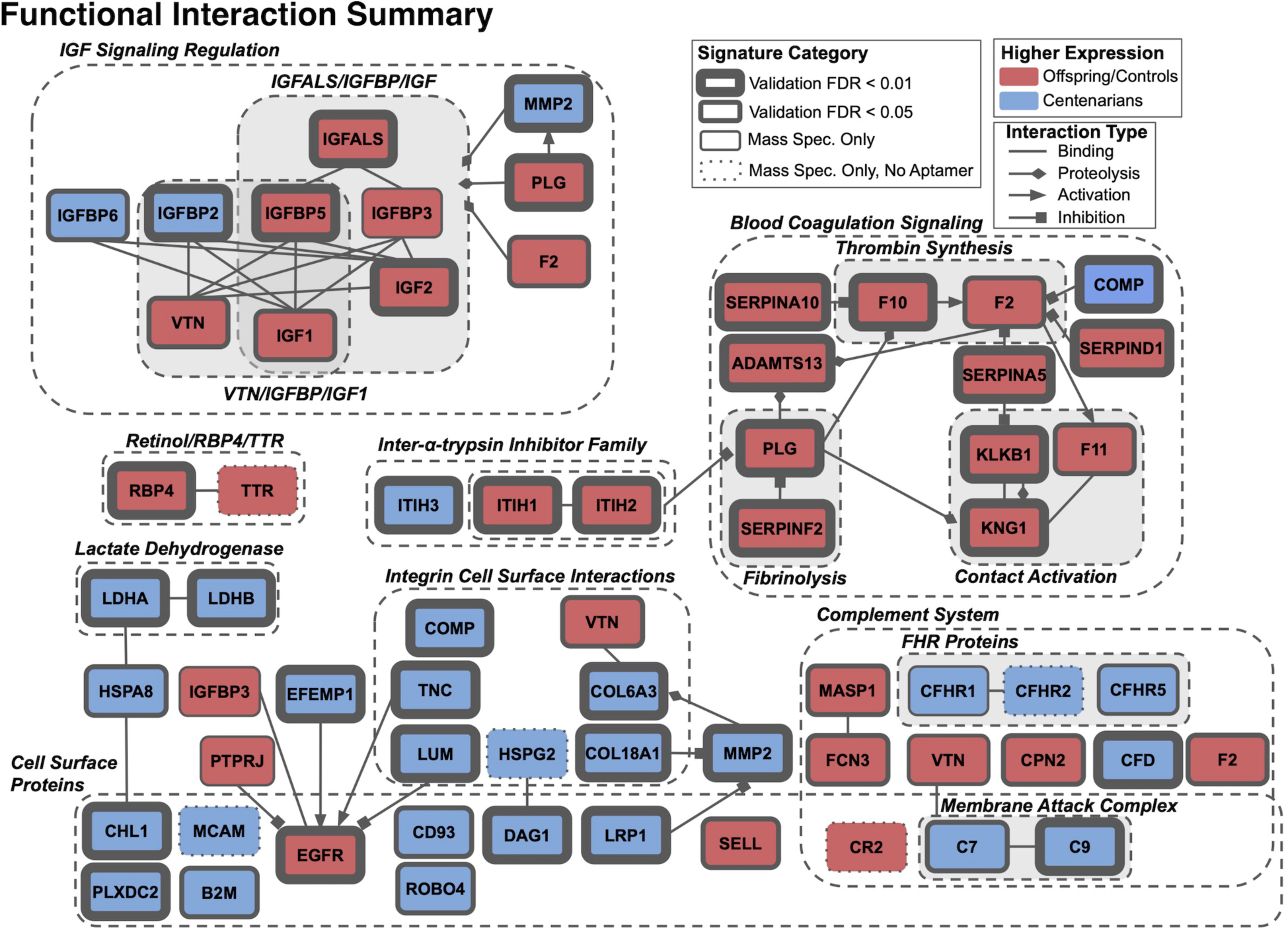
Summary of literature and Reactome database confirmed protein physical interactions. Additional information, including descriptions of interaction and sources can be found in Supplementary Table S5.

### Comparison with whole blood transcriptomic signatures of aging

We evaluated the concordance between our 80 protein, conserved serum proteomics signature of extreme old age at FDR < 0.05 and an independent whole blood transcriptomics signature of aging that we identified using 1346 expression profiles in the LLFS [16]. We identified this transcriptomic signature of age based on models with age as a continuous variable. Of 11,173 genes in the processed transcriptomics data, expression of 4,916 genes were associated with age at 1% FDR statistical significance threshold, with 2,570 and 2,346 genes associated with increased and decreased expression with age, respectively. Moreover, the 11,173 genes included 26 genes that encoded proteins included in the 80 conserved protein signature, of which 19 and 7 had higher expression in centenarians and offspring/control, respectively.

Of the 19 proteins with higher expression in centenarians, 12 coincided with genes with increased expression with age, and demonstrated significant over-representation as indicated by Fisher’s exact test, p-value = 0.0002. Alternatively, of the 7 genes with higher expression in offspring/control, only 1 coincided with genes with decreased expression with age, yielding non-significant over-representation, p-value = 0.808. The 12 gene/proteins demonstrating concordant higher expression levels associated with age, included ALDOA, B2M, CFD, CST3, LDHA, LRP1, LYZ, OAF, PLXDC2, PTGDS, S100A9, and YWHAZ. Functional annotation of these 12 proteins, yielded neutrophil degranulation (Reactome) as the only significantly enriched functional category, FDR = 0.036, which included six overlapping genes, ALDOA, B2M, CFD, CST3, LYZ, S100A9 (Figure 4).

**Figure 4:**
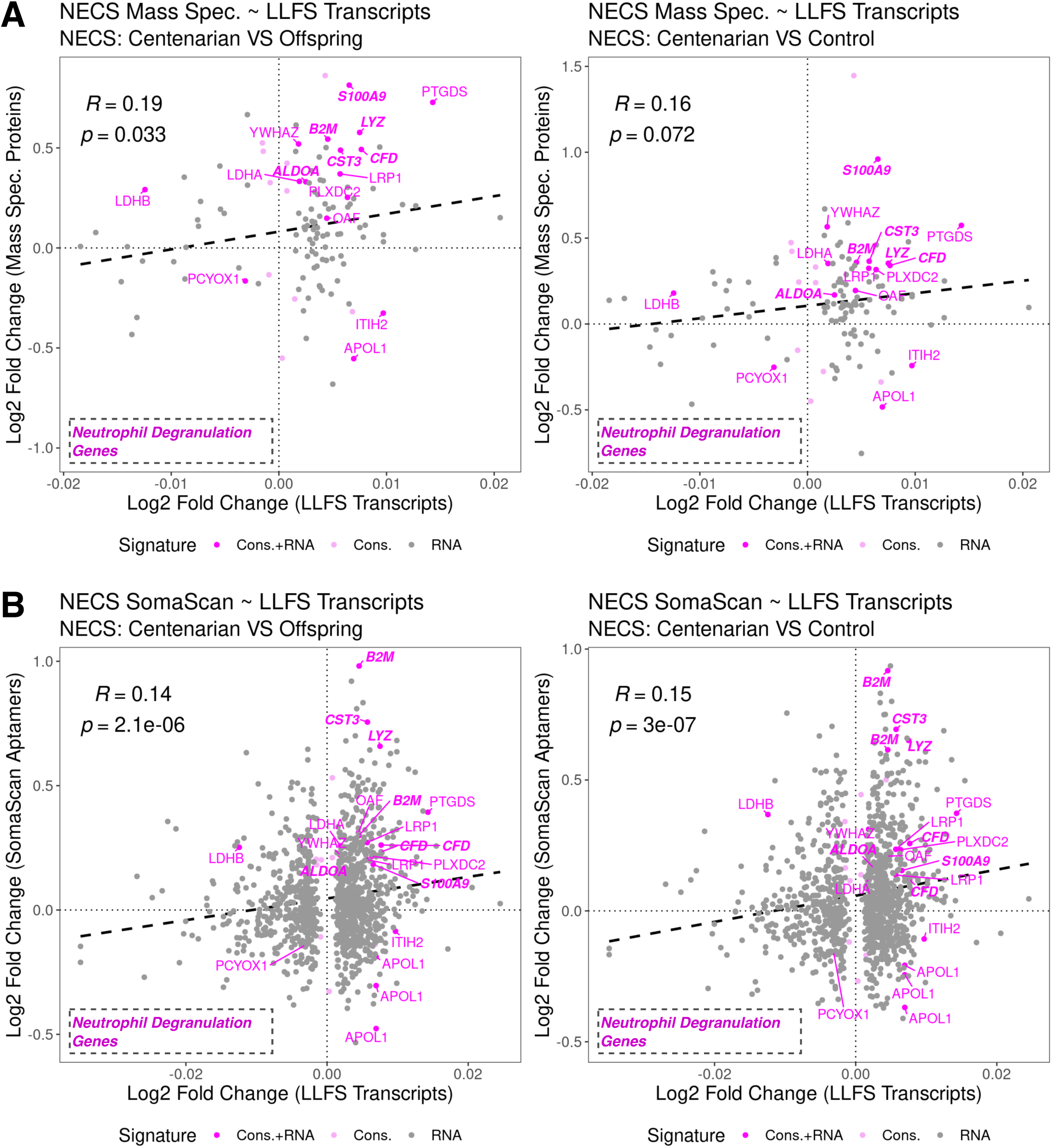
Comparison of mass spectrometry proteomic and LLFS transcriptomic results. Comparison of log2 fold changes from proteomic studies and LLFS gene expression models features characterized by either proteomic conservaRon analysis or transcriptomic analyses that are mappable across plaaorms. Proteomic results reflect differences between either offspring (led) or controls (right) to centenarian cohorts. LLFS transcriptomic results reflect age as a conRnuous variable, i.e. 1-year age differences. Proteins/genes annotated to the neutrophil degranulaRon pathway are highlighted in bold. A. Comparison of the mass spectrometry study and LLFS gene expression features. The plots comprise 123 coinciding protein:gene pairs. The full set of results for these features is reported in Supplemental Table S6. B. Comparison of the published SomaScan signature and LLFS gene expression features. The plots comprise 1,142 coinciding protein:gene pairs. The full set of results for these features is reported in Supplemental Table S7.

To evaluate the general concordances between the features identified by the transcriptomics study and the conserved protein signature, we evaluated the correlation between effect sizes across platforms for coinciding features of genes and proteins included in the 4,916 gene and 80 protein signatures. These included 123 and 1,142 coinciding features between the transcriptomic data and each of the mass spectrometry and published SomaScan studies, respectively. Since in proteomics studies the direction of effects was measured based on comparing younger cohorts to centenarians, we reversed this direction to make these results easily comparable to the transcriptomics signature. The full list of results for these features comparing the transcriptomic study compared to mass spectrometry and published SomaScan studies are reported in Supplemental Table S6 and Supplemental Table S7, respectively.

The transcriptomics data demonstrated general concordance with both proteomics studies based on trends of the direction and scale of age-associated effects (Figure 4). For these features, model coefficients comparing mass spectrometry and transcriptomic results were positively correlated across both mass spectrometry modeling results comparing centenarians to either offspring or controls (Figure 4A), yielding correlation estimates of 0.19 (p-value = 0.035) and 0.16 (p-value = 0.072), respectively. Comparisons between SomaScan and transcriptomic yielded complementary results (Figure 4B), with correlation estimates of 0.14 (p-value = 2.1E-6) and 0.15 (p-value = 3.0E-7), respectively.

## Discussion

Identifying robust serum proteomics signatures presents several challenges, particularly in terms of specificity, reproducibility, and interpretation of results. Although the SomaScan assay is highly regarded for its high coverage and reproducibility[33], non-specific aptamer-to-protein binding may lead to false positive results, thereby confounding biological inference [5]. On the other hand, mass spectrometry has comparatively low coverage and may exhibit higher technical variability when stringent criteria are imposed. Our cross-platform analyses of SomaScan and mass spectrometry assays sought to bridge their respective shortcomings and identify highly specific and robust proteomics signatures of extreme old age. In total, we characterized 80 proteins as being associated with extreme old age based on their conservation across the SomaScan and mass spectrometry studies at 5% FDR threshold. These 80 proteins comprised 23 proteins that were identified individually by both platforms, 38 proteins identified by the published SomaScan study only [15], 5 proteins identified by mass spectrometry only, and 14 proteins not identified by either study alone but reached significance based on inter-study conservation. Accordingly, these analyses confirm 61 proteins of the published SomaScan[15] study, and designate an additional 19 proteins with conserved association of extreme old age across the SomaScan and mass spectrometry studies (Table 2, Table 3). Finally, we characterized 16 additional proteins, which were only associated with extreme old age in the mass spectrometry study, five of which were measured by SomaScan, but were not identified as conserved across studies (Table 4).

These analyses yielded signatures of extreme old age that were highlighted by changes associated with numerous pathways, including blood coagulation, IGF signaling, extracellular matrix organization (ECM), and complement cascade. However, the interpretation of these results presents additional challenges stemming from general caveats of both proteomics and longevity studies. First, our proteomics profiles comprised only overall expression of proteins and did not probe the datasets for the evidence they may contain related to post-translational changes, which can impact expression quantification leading to confounded interpretation of differences between groups. Next, cross-sectional experiments comparing centenarians and younger cohorts fall short of explicitly identifying drivers of longevity, such that our results reflect changes related to both general and healthy aging, as well as the effect of being extremely old and close to the end of life. Accordingly, here we attempt to contextualize our results based on previous studies of aging, longevity, and disease.

### Blood Coagulation

Our analyses highlighted blood coagulation as an important process different in centenarians as compared to younger cohorts. The majority of proteins involved in blood coagulation were more abundant in younger cohorts. While, intuitively, this would suggest hypocoagulability in the centenarian cohort, previous studies have actually reported hypercoagulability in centenarians and aging. However, our findings most likely don’t contradict previous studies, but reflect that overall expression of a portion of these proteins may be negatively associated with their post-processing and activation. These proteins that were lower in centenarians compared to younger individuals included F2 (prothrombin/thrombin), F10, PLG (plasminogen/plasmin), and SERPINF2 (alpha 2-antiplasmin), KNG1 (high-molecular-weight kininogen (HMWK)/kinins), KLKB1 (prekallikrein/kallikrein), and F11. A previous study of coagulation markers from plasma of centenarians and younger cohorts reported significantly higher levels of prothrombin and coagulation factor X in younger controls[34]. However, these differences coincided with higher levels of coagulation activation markers in centenarians, including thrombin generation and F10 activation, as well as higher levels of the plasmin-antiplasmin complex in centenarians, indicative of elevated fibrin formation[34]. Moreover, plasma kallikrein cleaves HMWK to release bradykinin, which is composed of only nine amino acids [35]. Lower HMWK and higher kinin levels are associated with age-related diseases, including Alzheimer’s Disease (AD) and impaired cognitive function [36–39]. Finally, activated F11 is a component of both thrombin generation and the kallikrein-kinin system[40]. Taken together, it is likely our observed higher protein expression of coagulation markers in younger cohorts actually coincides with greater coagulation activity in the centenarian cohort.

In addition to SERPINF2, numerous serine protease inhibitors involved in blood coagulation were also more highly expressed in younger cohorts, including SERPINA10, SERPIND1, and SERPINA5, which collectively inhibit F10 [41], thrombin [42, 43], and kallikrein [44]. Moreover, SERPINA4, SERPINA6, and SERPINA10 were also more highly expressed in younger cohorts, while SERPINA11 was more highly expressed in centenarians. SERPINs have been previously reported as plasma markers of aging, including decreased SERPINF2 expression [45]. However, to our knowledge, associations of the other SERPINs with aging and longevity have not been previously reported.

### IGF signaling

Differences in the expression of proteins involved in IGF signaling regulation between centenarians and younger cohorts present a comprehensive depiction of this pathway that is consistent with previous studies of longevity and aging. First, all protein components of IGF/IGFBP/IGFALS ternary complex, including IGFALS, IGFBP3, IGFBP5, IGF2, and IGF1, were more highly expressed in younger cohorts, indicating that they have an elevated activity of this complex for regulation IGF1 and IGF2 activity. Other studies have reported that all of these proteins decrease with ages [46–49]. Alternatively, non-ternary complex components IGFBPs, IGFBP2 and IGFBP6 were more highly expressed in centenarians. Elevated IGFBP2 has been shown to be associated with AD and impaired cognitive function [50–52], and circulating IGFBP6 increases with age[53]. Moreover, our analyses identified three proteins involved in proteolysis of the IGF/IGFBP/IGFALS ternary complex, MMP2, PLG, and F2 [29], of which only MMP2 was more highly expressed in centenarians; this result also supports the findings of a previous study of long-lived individuals [54]. Considering that the assayed PLG and F2 expression may be negatively associated with their overall activity [34], it is likely that these observations reflect elevated IGF/IGFBP/IGFALS ternary complex proteolysis in centenarians, leading to the decreased levels of its components we observed in this study. Taken together, these results put forward a broad depiction of age associated IGF signaling regulation, harmonizing those reported by previous studies.

### ECM organization

Proteins that were more highly expressed in centenarians were mostly enriched for ECM organization processes. Aging is associated with reduced ECM integrity through collagen fragmentation and crosslinking, glycation, and accumulation of aggregation-prone peptides such as amyloid beta[55]. Our cross-platform analysis validated the higher expression of two collagens, COL18A1 and COL6A3, and the collagen degrading enzyme, MMP2 [56], in centenarians. In models of *C. elegans*, induced overexpression of collagens was shown to increase lifespan [57], however associations between COL18A1 and COL6A3 expression with human aging and longevity have not been previously detected. MMP2 has been shown to be elevated in individuals greater than 94 years old, but remains relatively stable throughout earlier life stages, suggesting a role in healthy aging [54]. Circulating endostatin produced by the cleavage of COL18A1 is positively associated with several age-associated diseases, including chronic obstructive pulmonary disease [58] and myocardial infarction [59]. Interestingly, endostatin inhibits MMP2 activity [60], suggesting that intact COL18A1 is a marker of healthy aging. Alternatively, in addition to the IGF/IGFBP/IGFALS ternary complex, MMP2 can cleave COL6A3 to generate endotrophin [61], which is associated with onset of obesity-related metabolic disorders [62]. However, MMP2 is more likely to capture intact COL6A3, and cannot characterize its cleavage activity. Taken together, these results illustrate the overall complexity of protein interactions involved in ECM organization and suggest a protective role of collagen activity for healthy aging in centenarians.

### Complement System

The complement system, which is part of innate immunity, was the only pathway to exhibit protein expression changes in both directions when comparing cohorts and centenarians. Complement system overactivation has been implicated in numerous age-associated diseases, including autoimmune and cognitive disorders [63]. We identified proteins involved in complement system activation, including some more abundant in centenarians and some more abundant in offspring/controls. However, these two groups of proteins generally comprise different components of this system. Proteins with higher expression in younger cohorts included lectin complement pathway components, MASP1 and FCN3, while protein more highly expressed in centenarians included alternative pathway components, CFD and alternative pathway complement factor H-related (FHR) proteins, CFHR1, CFHR2, and CFHR5, as well as membrane attack complex proteins, C7 and C9 [64, 65]. Given that the membrane attack complex is the endpoint of the complement system regardless of the initiating pathway, these results suggest an overall higher activation of the complement system in centenarians. Interestingly, previous studies have demonstrated opposite relationships of complement system activity for aging and longevity, however these reports have focused on complement factor, C3, which we did not identify. C3 is positively associated with age [66], but negatively associated with centenarian longevity [67]. Importantly, net C3 levels are not necessarily indicative of complement system activation, and are more precisely measured by the ratio between C3 and activated C3 [68]. Accordingly, these studies do not provide an explicit association between aging, longevity and overall complement system activation. Increased membrane attack complex levels in the choriocapillaris have been shown to increase with age and are associated with age-related macular degeneration (AMD) [69, 70].

In addition to broader pathways, our analysis revealed aging pathology-related subsets of proteins. Proteins more highly expressed in centenarians included lactate dehydrogenase components LDHA that did not reach a significant association in our original report and LDHB, which are marker for cell death and organ damage [71]. LDHA and LDHB are important enzymes of pyruvate metabolism and degrade pyruvate – a metabolite made from glucose through glycolysis into lactic acid. Higher levels of LDHA and LDHB in the centenarian cohort suggest dysregulation of glucose metabolism in extreme old age. Proteins more highly expressed in younger cohorts included retinol/RBP4/TTR complex proteins, RBP4 and TTR, of which circulating levels are negatively associated with insulin sensitivity [72, 73] and positively associated with cardiovascular events in elderly subjects [74, 75]. Accordingly, lower levels of RBP4 and TTR in the centenarian cohort indicate that they are potential markers of healthy aging.

Finally, the comparison between the serum protein signature and the whole blood transcriptomic signatures of aging from the LLFS revealed moderate concordance, highlighted by age-associated changes of six genes involved in neutrophil degranulation, ALDOA, B2M, CFD, CST3, LYZ, S100A9. Such moderate concordance in unsurprising, given the key differences between these studies, specifically that the serum proteomics studies primarily capture extracellular proteins in blood originating from a variety of tissues, while whole blood transcriptomics studies primarily capture intracellular transcripts in immune cells. Thus, consistencies between serum proteins and whole blood transcripts reflect specific incidences of transcripts from immune cells that are translated to proteins and eventually enter extracellular space. Accordingly, neutrophils are highly abundant in blood, comprising ∼70% of all immune cells [76], and neutrophil degranulation is a main component of their immune function [77]. Previous studies have shown age-associated changes in neutrophil function, including decreased chemotaxis and phagocytosis with age in adults throughout ages 30-79, but increased chemotaxis and phagocytosis in long-lived subjects, ages 90-103 [78]. However, explicit associations between neutrophil degranulation and aging have not been previously reported. Finally, these findings likely reflect age-associated shifts in immune cell composition, which we have previously characterized in a single-cell transcriptomics study of peripheral blood mononuclear cells (PBMCs) in centenarians and younger cohorts [32], although neutrophils have multi-lobulated nuclei and are not included in PBMC studies.

Our serum proteomics signatures of centenarians and younger cohorts strongly demonstrated changes to numerous pathways associated with health and aging. These results reflect the changes to both general and healthy aging, and likely more reflect the former. As a cross-platform study to identify conserved associations between SomaScan and mass spectrometry profiles, we stress the high-confidence of these signatures over those identified by single-platform studies. Accordingly, these results shed light on predominant mechanisms driving aging and longevity, thereby serving as roadmap for future studies to explore age-related pathology and possible interventions.

