## Supplemental Methods for "Cross-platform proteomics signatures of extreme old age"

**Supplementary Methods**

### Sample Preparation

*Reagents*: Pierce Quantitative Colorimetric Peptide kit, Pierce C18 spin columns, Pierce Top 12 Abundant Protein Depletion Spin Columns, TMT10plex Isobaric Label Reagent Set, TMT11-131C Label Reagent, UPLC grade acetonitrile, trifluoroacetic acid, and UPLC grade water were purchased from Thermo Fisher Scientific (Waltham, MA, USA). Trypsin/Lys-C Mix, Mass Spec Grade was obtained from Promega (Madison, WI, USA)., as well as the Amicon Ultra-0.5 mL Centrifugal Filters (3K NMWL), ammonium hydroxide solution (28-30% w/v), 2-chloroacetamide, ethyl acetate, and sodium deoxycholate, were purchased from Millipore Sigma (Burlington, MA, USA). 1M Tris, pH 8.5 was purchased from K D Medical (Columbia, MD, USA). LoBind 1.7 mL tubes were purchased from Eppendorf (Hamburg, Germany).

*Serum Protein Depletion, Processing, Proteolysis, and Labeling:* Serum samples were depleted of abundant proteins using the Pierce Top 12 Abundant Protein Depletion Spin Columns, according to the manufacturer's protocol. The remaining protein fraction was reduced, alkylated, and subjected to trypsin/LysC digestion, according to a previously published protocol [1]. A combined sample, composed of equal volumes of all participant serum samples, was used as a reference for all analyses. Briefly, an equal volume of each serum sample was depleted using Pierce Top 12 Abundant Protein Depletion Spin Columns (Thermo Fisher Scientific, Waltham, MA, USA), according to the manufacturer's instructions. Samples were concentrated using Amicon Ultra-0.5 mL Centrifugal Filters (3K NMWL, Millipore Sigma, Burlington, MA, USA) to ≤ 50 µL, by spinning at 14,000 RCF for 30 min. Next, 2X reduction and alkylation buffer was added to each sample to achieve a final concentration of 1% w/v sodium deoxycholate (SDC), 10 mM TCEP, 40 mM 2-chloroacetamide, and 100 mM Tris pH 8.5. Samples were heated for 10 min at 95 °C, then cooled to room temperature. Trypsin/LysC was added (1:100 enzyme-to-protein ratio), and proteolysis was performed at 37 °C for 4 hr. Peptides were acidified to a final concentration of 0.5% trifluoroacetic acid (TFA). Precipitated sodium deoxycholate was pelleted by briefly spinning samples (14000 RCF, 5 min). The supernatant was transferred to new tubes. Peptides were desalted based on Geyer PE *et al*., 2016 [2]. Pierce C18 spin columns were used for desalting; ethyl acetate with 1% TFA was used to remove sodium deoxycholate. Peptides were eluted from C18 spin columns in 80% acetonitrile, 19% dd$H_2O$, 1% ammonia, then dried under vacuum. The Pierce Quantitative Colorimetric Peptide Assay kit was utilized, followed by measurement of the absorbance at 480 nm on a Synergy HT microplate reader (BioTek, Winooski, VT, USA), to determine peptide concentrations. TMT labeling was performed according to the manufacturer's protocol. TMT reactions were quenched with 5% hydroxylamine, pooled, dried under vacuum, and resuspended in 500 µL 1% acetonitrile/99% water + 0.1% formic acid. Pooled peptide samples were desalted using a

SepPak C18 1cc cartridge (elution: 80% acetonitrile/20% water + 0.1% formic acid) and dried under vacuum.

*Peptide Fractionation:* To fractionate peptides, a high pH reversed phase separation was performed with $ddH_2O$, 1% acetonitrile, 0.1% $NH_4OH$ (buffer A) and $ddH_2O$, 80% acetonitrile, 0.1% $NH_4OH$ (buffer B). Peptide fractions were collected at 30 s. intervals from 5% B to 62.5% B over 30 min. The fractions were dried under vacuum, then suspended in 1% acetonitrile/99% water + 0.1% formic acid, for nUPLC-MS/MS.

*Liquid Chromatography and Mass Spectrometry:* nUPLC-MS/MS of peptides was performed on an Orbitrap Fusion Lumos Tribrid mass spectrometer (Thermo Scientific) with an online ACQUITY UPLC M-Class system (Waters) and a TriVersa NanoMate (Advion). A nanoEase Symmetry C18 UPLC Trap Column (100 Å, 5 µm, 180 µm × 20 mm, Waters) was used for peptide trapping. A nanoEase MZ HSS C18 T3 UPLC Column (100 Å, 1.8 µm, 75 µm × 100 mm, Waters) was used for peptide separation. Peptide trapping (4 µL/min, 4 min), was performed with 1% acetonitrile and 0.1% formic acid (Solvent A). Peptides were separated according to the following gradient: 0–1 min: 2% B, 1–3 min: 2–5% B, 3–43 min: 5–30% B (Solvent B: 99% acetonitrile and 0.1% formic acid). Conditions for MS analyses are as follows: positive mode, RF lens set to 30%. Orbitrap MS1 scans were acquired with 120,000 resolution (@ *m/z* 400), scan range *m/z* 400-1200, 1 µscan/MS, AGC target $2.0 \times 10^5$, and a 100 ms maximum injection time. Data-dependent acquisition was performed, with MS1 precursors filtered as follows: monoisotopic peak determination (MIPS) set to peptide mode, charge states 2-7, a 60 s. dynamic exclusion time after one fragmentation event, and a minimum intensity threshold of $3.0 \times 10^4$, with a maximum of eight MS2 scans for each cycle. HCD MS2 spectra were acquired using the following conditions: isolation in the quadrupole, isolation window of 0.7 *m/z.* HCD fragmentation (33% collision energy) was performed, and ions were analyzed in the Orbitrap (50,000 resolution @ *m/z* 400, scan range *m/z* 100–2000, 1 µscan/MS, AGC target $5 \times 10^4$, maximum injection time of 86 ms). Profile spectra were recorded. An additional fragmentation event (MS3) was performed using synchronous precursor selection, with 10 precursors selected from MS2, MS isolation window of 2.5 *m/z*, MS2 isolation window of 3.0 *m/z*, 55% collision energy, 50,000 resolution @ *m/z* 400, AGC target $5.0 \times 10^4$, 1 µscan/MS, detection in the Orbitrap, maximum injection time of 80 ms.

### Data Processing

MaxQuant was utilized to search and process TMT-labeled peptide data files, with the following settings: 10.0 ppm parent mass tolerance, 0.02 Da fragment mass tolerance, enzyme: Trypsin-LysC (cleavage at K, R), up to 3 missed cleavages, with TMT11plex (229.16) and carbamidomethylation (C) considered as fixed modifications. Reviewed protein sequences from the Uniprot *Homo sapiens* database*,* accession ID UP000005640, (downloaded Sept. 29,

2019), was used for all searches. Protein matches were filtered based on a 1% false discovery rate threshold, and ≥ 1 unique peptide.
